# Identification of genetic variants affecting vitamin D receptor binding and associations with autoimmune disease

**DOI:** 10.1101/080143

**Authors:** Giuseppe Gallone, Wilfried Haerty, Giulio Disanto, Sreeram Ramagopalan, Chris P. Ponting, Antonio J. Berlanga-Taylor

**Author notes:** Authors contributed equally (Sreeram Ramagopalan, Chris P. Ponting, Antonio J. Berlanga-Taylor).

## Abstract

Large numbers of statistically significant associations between sentinel SNPs and case-control status have been replicated by genome-wide association studies. Nevertheless, few underlying molecular mechanisms of complex disease are currently known. We investigated whether variation in binding of a transcription factor, the vitamin D receptor (VDR) whose activating ligand vitamin D has been proposed as a modifiable factor in multiple disorders, could explain any of these associations. VDR modifies gene expression by binding DNA as a heterodimer with the Retinoid X receptor (RXR).

We identified 43,332 genetic variants significantly associated with altered VDR binding affinity (VDR-BVs) using a high-resolution (ChIP-exo) genome-wide analysis of 27 HapMap lymphoblastoid cell lines. VDR-BVs are enriched in consensus RXR::VDR binding motifs, yet most fell outside of these motifs, implying that genetic variation often affects binding affinity only indirectly. Finally, we compared 341 VDR-BVs replicating by position in multiple individuals against background sets of variants lying within VDR-binding regions that had been matched in allele frequency and were independent with respect to linkage disequilibrium. In this stringent test, these replicated VDR-BVs were significantly (*q* < 0.1) and substantially (> 2-fold) enriched in genomic intervals associated with autoimmune and other diseases, including inflammatory bowel disease, Crohn’s disease and rheumatoid arthritis. The approach’s validity is underscored by RXR::VDR motif sequence being predictive of binding strength and being evolutionarily constrained.

Our findings are consistent with altered RXR::VDR binding contributing to immunity-related diseases. Replicated VDR-BVs associated with these disorders could represent causal disease risk alleles whose effect may be modifiable by vitamin D levels.

## Introduction

Genetic variants can alter transcription factor (TF) binding, chromatin status and gene expression [1, 2, 3] and can be explanatory of disease susceptibility variation. Rather than residing in protein-coding sequence, the majority (approximately 93%) of disease-or trait-associated variants lie in non-coding sequence [4]. Furthermore, non-coding variants in regulatory elements explain 8-fold more heritability of complex traits than protein-coding variants [5]. If we are to better understand the mechanistic bases to complex disease (and their interaction with environmental factors) then we will need to understand how sequence differences in TF binding sites (TFBSs) across multiple genotypes contribute to disease susceptibility. Variations in TF affinities will then need to be cross-referenced not just with the sentinel variant from genome-wide association studies (GWASs), which has a very low probability (∼5%) of being causal and is on average 14 kb from the true causal variant [6], but also with all other variants with which it lies in strong linkage disequilibrium (LD). True causal variants will be enriched in *cis*-regulatory elements, especially in enhancers and, to a lesser extent, in gene promoters [6]. These elements are often active in only a few cell types [7] and at limited developmental time points [8], which provides an opportunity for relating genetic variation — via molecular observations (TF binding differences) — to cellular or organ-based aetiologies.

Understanding complex disease will also need the determination of how functional genetic variants interact with environmental factors. One such factor is Vitamin D, a class of fat-soluble secosteroids that enhance intestinal absorption of dietary minerals, which are synthesised in the skin upon exposure to sunlight. Many observational studies have yielded associations between low serum concentrations of 25-hydroxyvitamin D [25(OH)D] and the risks of developing diverse diseases [9, 10]. However, Mendelian randomisation studies, for example on susceptibility to multiple sclerosis [11] and all cause mortality [12] have indicated a causal role for this hormone. Identifying a molecular basis to these statistical associations would also indicate a direct causal role for vitamin D levels in these diseases. Vitamin D signalling occurs principally following the binding of calcitriol, the active form of vitamin D, to its cognate nuclear vitamin D receptor (VDR) which binds DNA in a heterodimer with the Retinoid X receptor (RXR [13]). We previously have shown that VDR binding in lymphoblastoid cell lines (LCLs) occurs at 2,776 locations and preferentially (> 2-fold) within intervals genetically associated with diseases, such as Crohn’s disease and Multiple Sclerosis [14]. On one hand, these enrichments could indicate a direct modifying effect of VDR-binding on disease risk. On the other hand, they could, more trivially, reflect a general association between regulatory regions and disease-associated genomic intervals. These enrichments of VDR binding within these intervals also are not informative of whether altered VDR binding to DNA functional elements explains genetic contributions to disease risk. If this is the case, we then also wish to determine whether it is a gain or loss of VDR binding that causally increases disease risk.

In this study, we inferred VDR-binding sites using ChIP-exo (chromatin immunoprecipitation combined with lambda exonuclease digestion followed by high-throughput sequencing [15, 16]) from each of 27 LCL samples using 3 complementary approaches. The first ‘peak-calling’ method models the variation in read density. The second and third approaches predict VDR-binding sites using genetic variation to explain differences in VDR-binding affinity, either by quantitative trait loci association testing (‘QTL’) across all samples or by allele-specific binding (‘ASB’) analysis of allelic imbalance based on read depth at sites with heterozygous single nucleotide variants. Sequence variants were identified that both alter VDR binding and have been statistically associated by GWAS with altered risk for particular diseases (or are in strong LD with such variants). These are excellent candidates for sequence variants that, through their alteration of VDR binding, directly alter disease susceptibility. We considered whether VDR-binding events have consequences on fitness, and thus when lost or gained either provide protection from or confer additional risk of disease. Compared against a background of all VDR-binding sites, we found that human variants associated with variable VDR-binding are enriched (by up to 2-fold) in genomic intervals previously associated with particular traits, including some autoimmune diseases.

## Results

### ChIP-exo yields finely resolved VDR binding peaks genome-wide

Calcitriol–stimulated lymphoblastoid cells for 30 HapMap samples were grown and prepared for ChIP essentially as described previously [14] with modifications for ChIP-exo [15, 16] (Methods; Supplementary Table 1). In total, 27 samples successfully passed quality checking (Supplementary Material). Unlike for data from the more traditional ChIP-seq approach, analysis of ChIP-exo data has yet to become standardised. We thus undertook an in-depth study using contrasting approaches to modelling the ChIP-exo data background, to handling duplicate reads, to performing cross-correlation analyses and to calling peaks; these approaches are discussed in detail in the Supplementary Material (see also Supplementary Figures 1 and 2, Supplementary Tables 2–9).

VDR peaks were both highly reproducible between samples, with 15,509 intervals containing peaks called in at least 3 samples (CP_*o*3_ peak set) (Figure 1A and B; Supplementary Figures 3 and 4), and highly concordant with our previously published VDR ChIP-seq data for calcitriol-stimulated LCLs (CP_*o*3_: 76.3%; Figure 1C; Supplementary Table 11) [14] (Supplementary Material). For peaks identified in both experiments, binding intervals were considerably better resolved (typically > 5-fold) using ChIP-exo (Supplementary Figure 2A, pairwise Kruskal-Wallis Test, *p* < 0.001).

**Figure 1:**
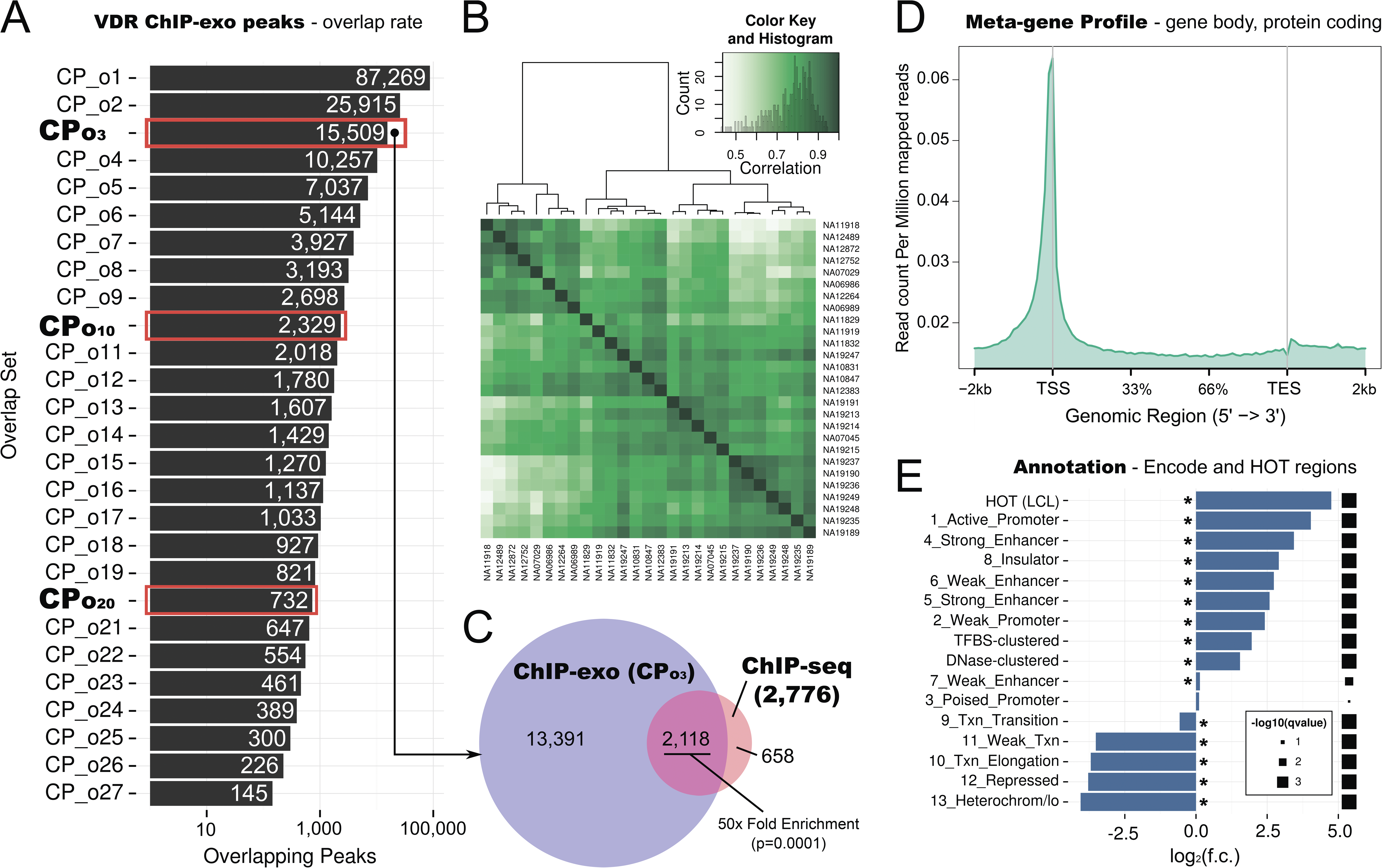
ChIP-exo peaks are reproducible among samples and with previous ChIP-seq results, and are enriched in promoters and enhancers. **A:** Numbers of overlapping ChIP-exo peaks across 27 LCL samples (log scale). Highlighted rows indicate numbers of peaks called in at least three (CP_*o*3_), ten (CP_*o*10_) or twenty (CP_*o*20_) samples. **B:** Normalised peak read depths [63, 62, 78] (Supplementary Material) are concordant between samples. VDR ChIP-exo peak binding affinity heatmap, showing pair-wise binding affinity correlation among all 27 sample pairs. (*Inset*) histogram of correlation counts. **C:** 50-fold enriched overlap between CP_*o*3_ peaks and the consensus ChIP-seq peakset from [14] (enrichment test: GAT [64], Supplementary Material). **D:** Meta-gene profile of VDR binding reads. The panel shows a meta-profile of read pile-up for the pooled 27 LCL samples, averaged over gene bodies (Ensembl v. 75, protein coding genes). **E:** Enrichment of VDR binding sites in the CP_*o*3_ consensus with ENCODE chromatin states (chromHMM [79]) from LCL NA12878, TF-dense regions (HOT [17] and ENCODE clustered TF binding sites) and DNase I hypersensitive areas (ENCODE). Asterisk denotes significance at 1% False Discovery Rate (FDR) threshold (enrichment test: GAT [64], Supplementary Material).

**Figure 2:**
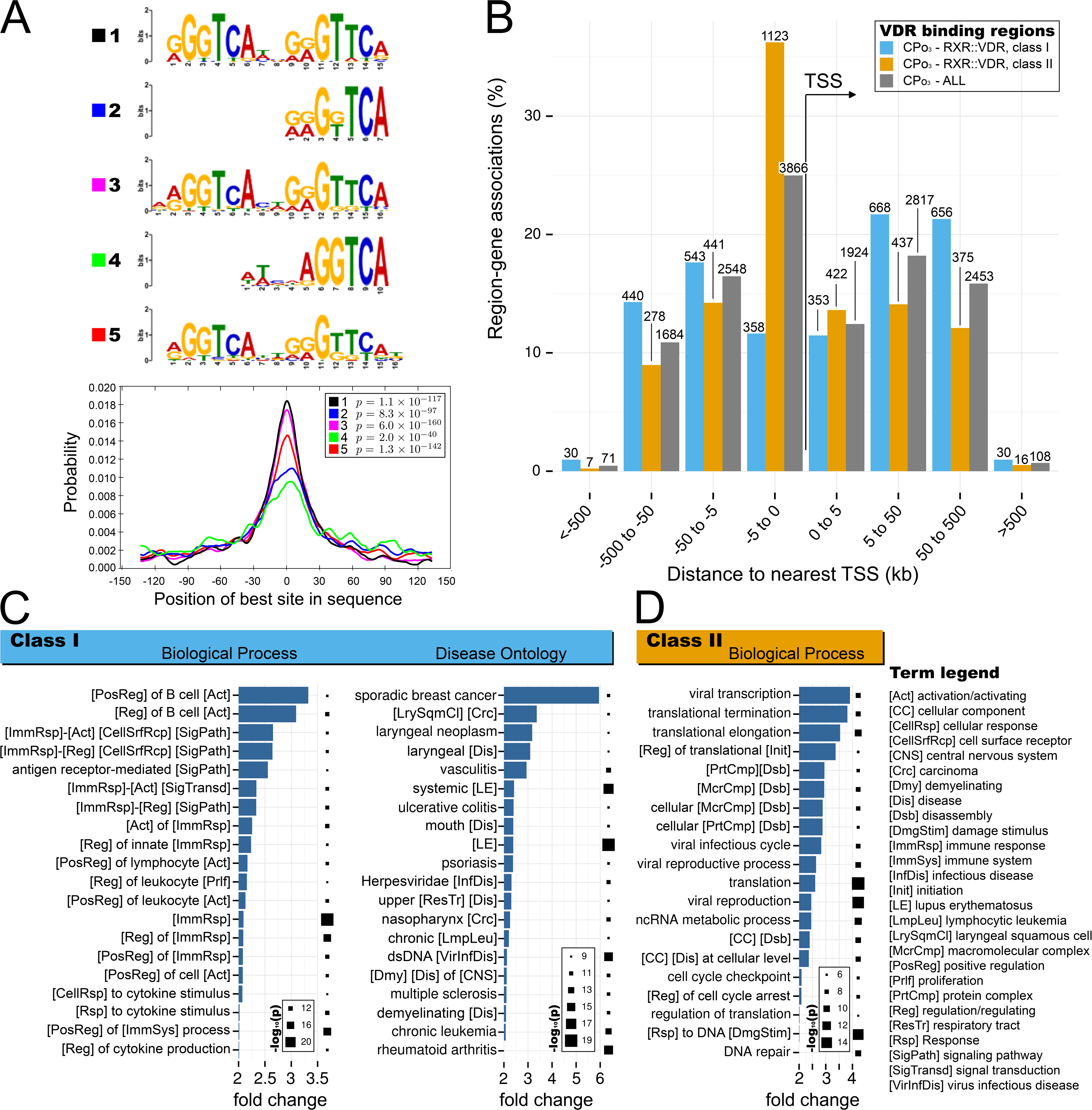
VDR binding intervals are immune-associated or are housekeeping-associated tending to be either present far from, or adjacent to, transcriptional start sites, respectively. **A:** The most highly ranking motif cluster ensuing from a *de novo* motif analysis of the VDR CP_*o*3_ consensus intervals using MEME-ChIP [65], including a CentriMO [80] analysis of enrichment with respect to VDR peak centres. B-D: GREAT analyses [81] for subsets of VDR peaks from the CP_*o*3_ consensus set containing either a strong-to-perfect (class *I*) or weak-to-non-existent (class *II*) instance of the RXR::VDR heterodimer motif. **B:** Distributions of distances between known TSS sites (UCSC hg19, Feb 2009) and VDR binding locations for classes I or II, or all, VDR peaks. **C,D:** Enrichment for Gene Ontology Biological Process and Disease Ontology terms for class *I* (**C**) or class *II* (**D**) RXR::VDR containing sites, respectively. For **C** and **D**, bars map to fold change whereas Hinton plots map to −log_10_ of the enrichment significance *p*-value.

As expected, VDR binding peaks occurred preferentially at protein coding transcriptional start sites (TSS) (Figure 1D) and within regions of open chromatin (at LCL DNase 1 hypersensitivity sites, DHS), especially for those observed repeatedly (Supplementary Figures 5 and 6). VDR binding peaks were significantly concentrated within proposed functional elements, specifically regions of high regulatory factor binding (HOT regions [17] and clustered TF binding sites from ENCODE), active promoters, enhancers and insulators (Figure 1E; Supplementary Figure 7) and previously reported GWAS disease loci, principally for autoimmune disorders (Supplementary Figure 8). Notably, this latter observation could reflect a causal effect in which VDR-binding influences disease susceptibility and/or with gene loci being correlated non-causally with both disease susceptibility intervals and VDR binding sites.

### VDR binding motifs occur preferentially in enhancers

*De novo* motif analyses of peaks within ChIP-exo VDR binding intervals revealed motifs strongly resembling those for the RXR::VDR DR3 heterodimer [18, 19] (Figure 2A; Supplementary Material, Supplementary Figure 9A and Supplementary Table 12). For example, 9,899 (63.8% of the CP_*o*3_ set) instances of this DR3 heterodimer motif occurred in binding sites (PScanChIP [20], score > 0.7). 874 instances of the monomeric VDR motif were also identified in these peak intervals (Supplementary Figures 9B; PScanChIP score = 1) which are known to have limited affinity for functional dimeric nuclear hormone receptor complexes [18]. The heterodimeric DR3 motif was strongly over-represented (Supplementary Table 12) both locally (in the peak sequences, compared to neighbouring regions) and globally (in the peak regions, compared to LCL DHS sites [20]) and was significantly centrally enriched in the binding regions (Figure 2A).

VDR binding sites could be separated into functionally distinct classes based on the presence of RXR::VDR DR3-like motifs. *Class I* sites contain strong DR3-like motifs (top 20% of the PScanChIP score distribution, 3,094 regions, Supplementary Figure 10A) and tend to be located away from genes’ transcriptional start sites (TSSs) and yet close to genes involved in immune response processes (Figure 2B,C). *Class I* VDR binding sites show greatest significance of proximity to genomic regions associated with cancer and autoimmune diseases (Figure 2C). We compare these to *Class II* sites which contain only weak or no DR3-like motifs (bottom 20% of PScanChip scores, 3,102 regions, Supplementary Figure 10A) (Figure 2B,D). Class *I* binding events occur preferentially in strong or weak enhancers defined by ENCODE, rather than in promoters (Supplementary Figure 10B and 10C).

### Thousands of variants explain differential VDR binding

We next used the QTL and ASB methods to explain differences in VDR binding affinity (Supplementary Material). The QTL analysis used Bayesian regression modelling of variants underlying binding peaks [21, 22] (Supplementary Material, Supplementary Tables 13 and 14, Supplementary Figures 12 to 15). The ASB analysis of allelic imbalance is based on read depth at heterozygous single-nucleotide variants [23] and uses sample-specific maternal and paternal reference maps, which we corrected for unannotated copy number variation and from which we excluded variants in ENCODE blacklisted repeat regions [24]. The two methods’ variants associated with differences in VDR binding (VDR-Binding Variants, VDR-BVs) were annotated and prioritised based on the guidelines proposed by [25] and related algorithms (Funseq2, [26]).

We report a total of 43,332 VDR-BVs. Of these, only 6.6%-10.5% (2,867 or 4,447) lie within (or ‘hit’) RXR::VDR consensus motifs (Pscanchip score > 0.7 or > 0.5, respectively). This could be explained, in part, by variants that alter DNA-binding affinity either for other subunits of a larger multi-molecular complex (‘collaborative binding’ [27]) or for factors that inhibit RXR::VDR binding.

Consequently, we next considered whether sequence variation in motifs for potential binding cofactors of RXR::VDR could explain the remaining > 90 % of binding variation. We found that position weight matrices for 26 additional TFs (Supplementary Tables 15 and 16) were significantly enriched (*p* < 0.1) (i) relative to both open chromatin genome-wide and chromosomally adjacent regions (< 150 bp), and/or (ii) at the peak summits within VDR-BV intervals (Pscanchip score > 0.5 or 0.7; Supplementary Methods). Virtually all (84.1-88.3%) of binding variation could be explained by VDR-BVs lying within a RXR::VDR consensus motif, or within one or more of these 26 additional motifs (Pscanchip score > 0.7 or > 0.5). These are upper-bound values because of the inaccuracy of motif prediction and because not all variants lying in motifs will alter binding affinity.

### Functional annotation of VDR-BVs

To begin to understand whether VDR-BVs could contribute to particular human diseases, we first considered their enrichment in particular genomic annotations. For this, we were interested in whether these variably-bound sites are enriched relative to a background of all VDR binding regions genome-wide so that we account for the non-uniform chromosomal distribution of binding sites. We chose for our null expectation all variants that were tested for differential binding within replicated (CP_*o*3_) VDR binding peaks (Supplementary Material). Relative to this stringent background, VDR-BVs are 20%-40% enriched within enhancers but not in promoters (Figure 3A). These patterns of enrichment are reinforced when we stringently select only the most significant VDR-BVs (VDR-sBVs, Supplementary Figure 16A; AlleleSeq FDR ≤ 0.01; 6,715 VDR-BVs) or the 357 recurrent VDR-BVs that are called at the same position in multiple samples (VDR-rBVs, Supplementary Figure 16D). We conclude that VDR-BVs are not uniformly distributed among all VDR binding genomic intervals, and are especially frequent in enhancer regions manifested in LCLs.

**Figure 3:**
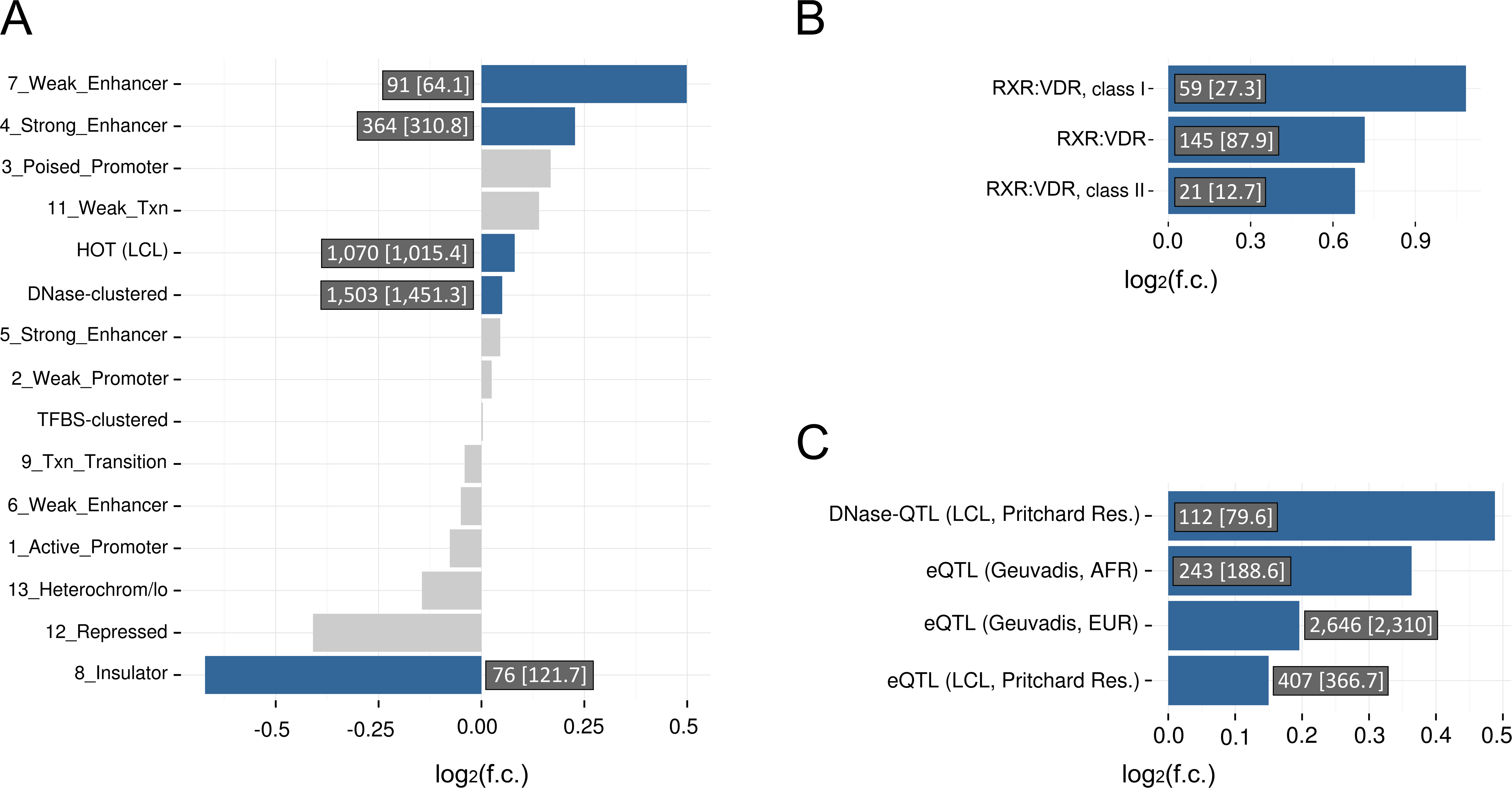
VDR-BVs are enriched at enhancers, in RXR::VDR DR3-type motifs, and in LCL eQTLs. Genomic association testing of VDR-BVs with functional annotation. **A:** Enrichment of VDR-BVs within ENCODE chromatin state segmentation tracks for LCL NA12878, TF-dense regions (HOT [17] and ENCODE clustered TF binding sites) and DNase hypersensitive areas (ENCODE). **B:** Enrichment of VDR-BVs in VDR consensus motif intervals present in VDR CP_*o*3_ binding regions. **C:** Enrichment of VDR-BVs at DNase-QTLs and eQTLs from the Pritchard resource and CEU/YRI LCL eQTLs from the GEUVADIS resource. Significant enrichments are indicated using blue histogram bars (f.c. = fold change of observed versus expected overlaps; FDR = *q* < 0.1; numbers in a box close to each bar indicate ‘observed [expected]’ overlap counts; grey bars indicate lack of significance). For panel **A**, enrichments shown are above-and-beyond the previously observed (Figure 1E) enrichments of VDR CP_*o*3_ peaks in the same functional annotation classes (relative to a background of 20,330 1000 Genomes SNPs in CP_*o*3_ VDR binding regions). For panel **B**, enrichments are relative to a background of 114,155 1000 Genomes variants lying under VDR ChIP-exo read pileups (≥ 5 reads) which had been tested as potential VDR binding affinity modifiers. This background was further corrected for the analyses in panel **C**, were only LD-independent foreground VDR-BVs were tested and 10,000 DAF-matched random background sets were extracted with replacement from the main set of 114,155 background variants.

VDR-BVs occurred 60% more frequently in RXR::VDR motifs within replicated (CP_*o*3_) VDR binding peaks and 120% more frequently in strong (class I) RXR::VDR motifs in the same regions (Figure 3B). Again, these enrichments strengthened substantially for VDR-sBVs or VDR-rBVs (Supplementary Figure 16B and 16E, respectively).

We then considered whether VDR-BVs are significantly enriched within LCL expression QTLs (eQTLs) relative to 1000 Genomes variants under VDR ChIP-exo pileups tested for VDR binding variation potential (Supplementary Material). In addition, these background variants were matched by derived allele frequency and LD-dependence confounders (Methods). We also took care to only consider variants that do not share strong linkage disequilibrium (32,183 VDR-BVs, 5,808 VDR-sBVs and 341 VDR-rBVs.)

VDR-BVs were enriched with dsQTLs (1.4-fold) and eQTLs (1.1-1.2-fold; Figure 3C) particularly for the more stringent VDR-BV subsets (Supplementary Figures 16C and 16F). The enrichment in eQTLs was confirmed using an independent eQTL resource (GEUVADIS LCL eQTLs [28]; Supplementary Material).

### Complex trait-associated variants influence VDR binding

To investigate whether susceptibility to specific diseases could be influenced by VDR-BVs, we considered their locations relative to GWAS variants significantly associated with 472 diverse diseases or traits from 688 studies in the largest catalogue available (GRASP v2.0 [29]) with at least 5 associated genome-wide significant SNPs (*p* < 5 × 10^−8^) (Supplementary Material). All traits available in these catalogues were considered so as not to bias subsequent findings. Again, we employed stringent procedures, matching variants for allele frequency and LD-dependence and discarding VDR-BVs in the MHC region as well as again comparing against a background of all VDR-binding regions (“Test 3”, p. 34 of the Supplementary Material).

VDR-BVs were significantly and up to 2-fold enriched within LD intervals associated with 17 diseases or traits (Figure 4). It is notable that these are not drawn randomly from all 472 traits considered, but are mostly immune-or inflammatory-related disorders, such as sarcoidosis, Graves’ disease, Crohn’s disease, irritable bowel syndrome and type I diabetes (Figure 4A). The most enriched trait is Alzheimer’s disease, for which 25-hydroxyvitamin D level is a proposed causal risk factor [30].

**Figure 4:**
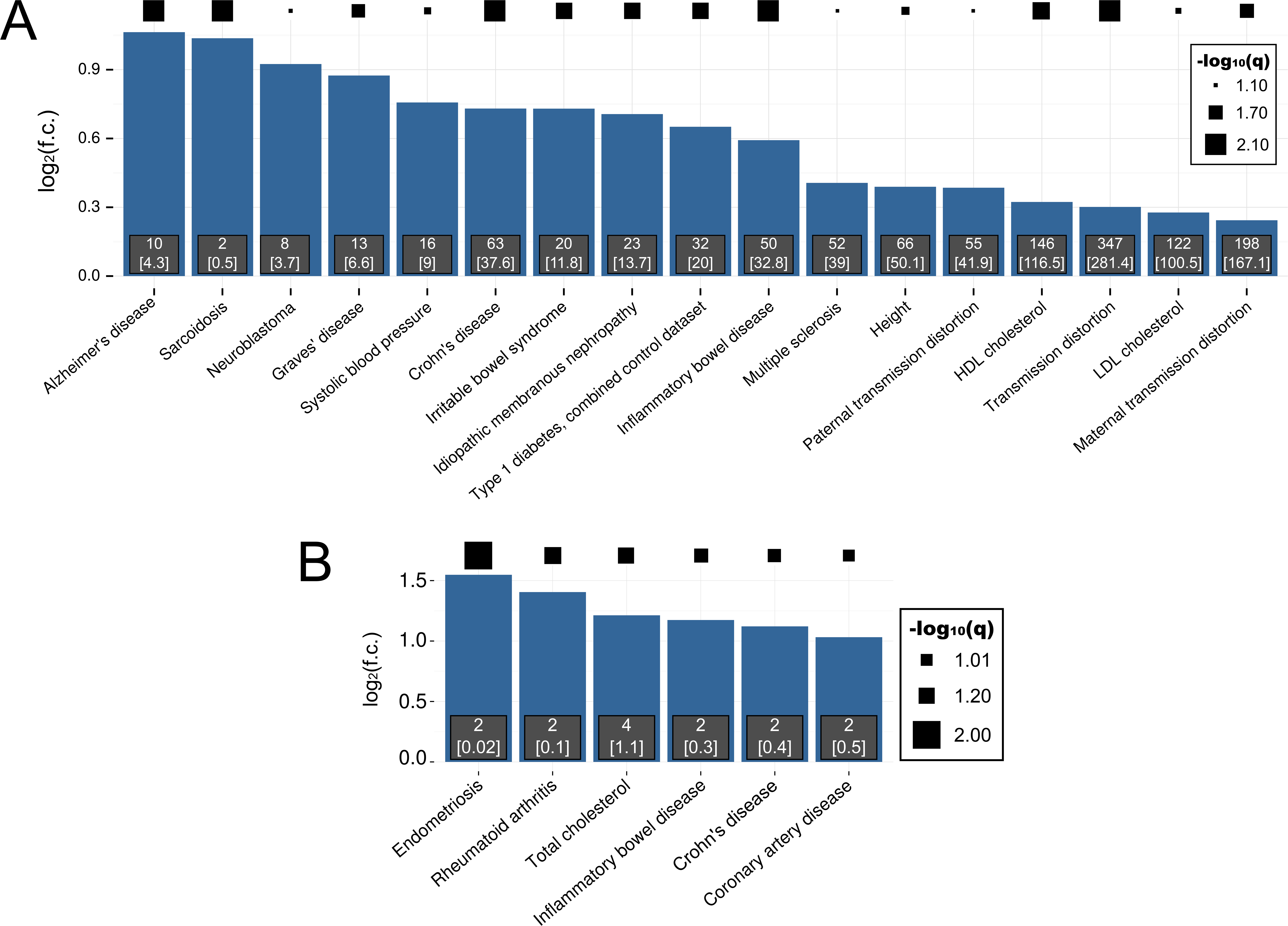
VDR-BVs are enriched in GWAS LD blocks for autoimmune, inflammatory and other diseases. **A-B:** Traits and diseases showing enrichment of VDR-BVs (**A**) or VDR-rBVs (**B**). Diseases/trait associations were acquired from the GRASP catalog v2.0 [29]. Statistical association was computed between VDR-BVs and strong-LD (Supplementary Material) intervals around genome-wide significant (*p* < 5 × 10^−8^) disease tag-SNPs from the GRASP catalog. All available GRASP diseases and traits represented by at least 5 SNPs were analysed and those showing significant enrichment of VDR binding beyond a 1% FDR threshold were retained; disease associations supported by only 1 VDR-BV were discarded (full data are presented in the Supplementary Material). The sizes of black squares indicate the statistical significance of enrichments.

For the 341 VDR-rBVs, intervals from six disorders contained more VDR-rBVs than expected from sets of DAF-matched LD-accounted regions that bind VDR (Figure 4B). We emphasise that this represents a highly stringent analysis that accounts for all known confounding effects, namely potential biases from called VDR-binding sites, population stratification and physical linkage.

### Functional impact of genetic variation on loss or gain of VDR binding

This evidence is consistent with altered RXR::VDR binding contributing to immunity-related diseases. Nevertheless, because variants in TFBS often fail to alter binding affinity and/or have no measurable effect on gene expression levels [31, 32], we needed to demonstrate that VDR-BVs are functional, specifically that they influence gene expression and have been negatively selected over evolution [33]. As expected if they are enriched in functional variants, we found that 50%-100% more VDR-BVs than random samples lie near (< 10 kb) genes that are differentially expressed, in LCLs, upon addition of calcitriol [14] (*p* < 10^−4^, Supplementary Methods).

Also, as expected if as a class they are functional, replicated VDR-binding peaks exhibit variable cross-species conservation [34] across RXR::VDR motif positions (Supplementary Figure 17A and B). This signature of uneven selection across the DR3 motif is evident for class I VDR binding sites (Supplementary Figure 17C and D; Kruskal Wallis test, *p* = 1.9 × 10^−11^ and *post-hoc* pairwise Kruskal Wallis comparisons), but not in class II sites (Kruskal Wallis test, *p* = 0.14) consistent with the DR3 heterodimer-binding motif regulating transcription.

Next we assessed the impact on VDR binding affinity of VDR-BVs with respect to their location within the consensus DR3-type motif (JASPAR database, Figure 5), the transcriptionally active binding site motif for the RXR::VDR heterodimer [35, 36, 37]. Mapping of allele-specific ChIP-exo reads to multiply replicated VDR binding regions (CP_*o*3_ peaks) was highly predictive of whether variants substantially alter (‘break’) functional motifs. For 89% of observations, the change in the strength of the VDR binding motif correctly predicts the direction of VDR binding affinity change (Figure 5A), a proportion that rises to 100% in class I VDR binding sites (Supplementary Figure 18A). Motif breaks do not favour 5’ RXR over 3’ VDR hexamers (Figure 5A, Wilcoxon rank sum test, *p* = 0.71), but especially impacts conserved G and C nucleotides (positions 2, 5, 11 and 14), and a T nucleotide (position 13). Notably fewer events occur at the near-essential T nucleotides at positions 4 and 12. Interestingly, in class I sites there are no VDR-BVs disrupting the T nucleotide at position 12 (Supplementary Figures 18A) which could reflect either low mutation rates (mutational ‘cold spots’) or strong purifying selection against deleterious nucleotide substitutions at these binding sites [38].

**Figure 5:**
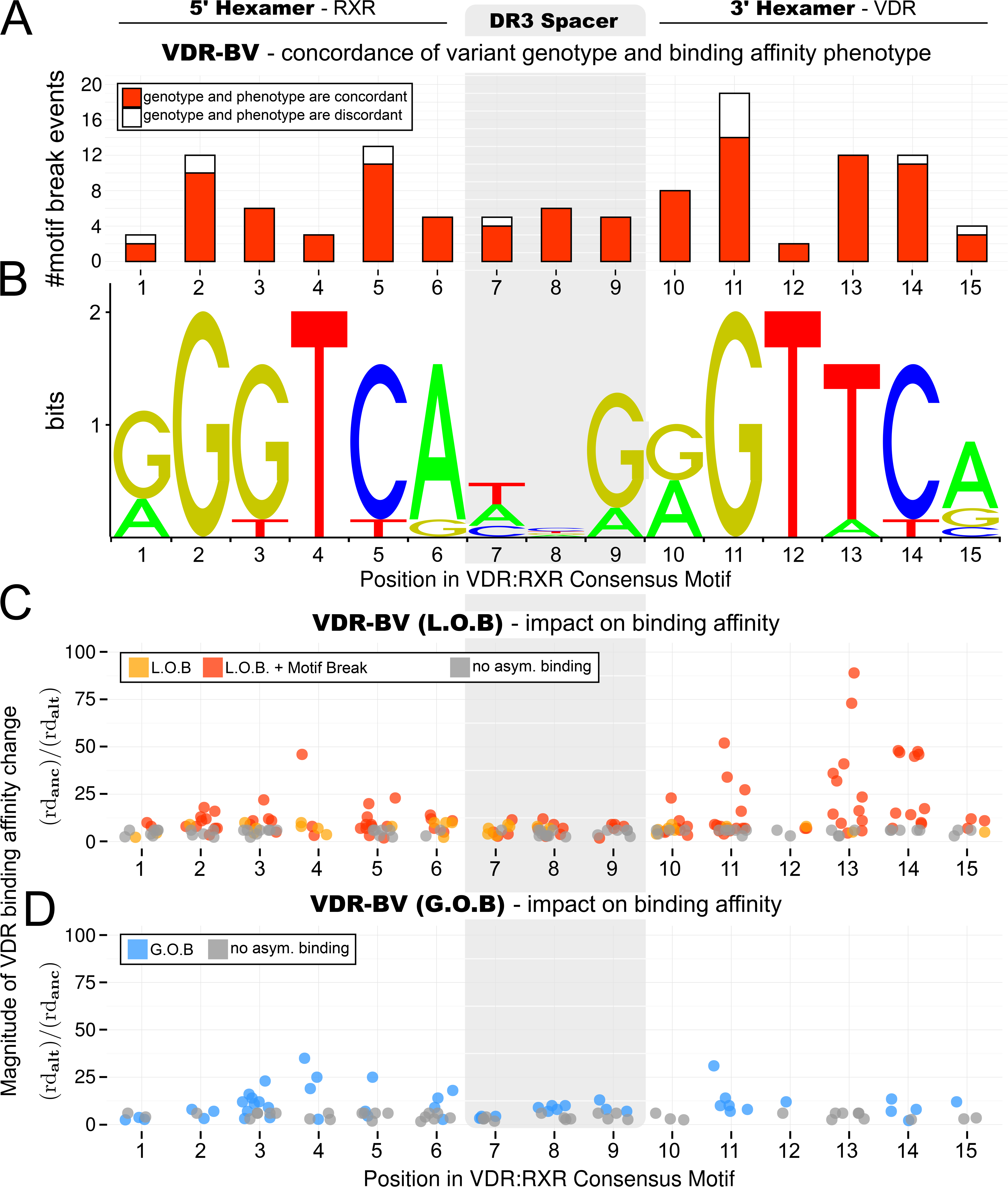
Genetic variation modulating VDR binding affinity in LCLs within the canonical RXR::VDR heterodimeric binding motif. Genome-wide quantification of the effect of genetic variation on VDR binding, based on the analysis of VDR-BVs in CP_*o*3_ binding peaks which hit the canonical RXR::VDR DR3 heterodimeric consensus motif (motif shown in panel **B**, JASPAR database, [19]). **A:** Distribution across the RXR::VDR motif of VDR-BVs which cause significant motif disruption (significance assessed via FunSeq2 using TFMPvalue [82], with default threshold *p* < 4 × 10^−8^) and quantification of the concordance between directionality of VDR PWM disruption and directionality of resulting VDR binding affinity variation. 102/115 (89%) VDR-BVs in CP_*o*3_ VDR peaks that significantly break the RXR::VDR motif predict the direction of VDR binding affinity change. **C,D:** Genome-wide quantification of the phenotypic effect of all VDR-BVs intersecting the RXR::VDR motif (including those which do not generate a motif break at the above significance level). The vertical axis (*Magnitude of VDR binding affinity change*) indicates the fold change of read depth of the ancestral versus the derived allele (panel **C**) or derived versus the ancestral allele (panel **D**). **C:** Impact on VDR binding affinity of Loss of Binding (LOB, orange dots) VDR-BVs and Loss of Binding VDR-BVs which test for significant RXR::VDR motif break (red dots). **D:** Impact on VDR binding affinity of Gain of Binding (GOB) VDR-BVs (blue dots). Nucleotide position has a significant influence on LOB (Kruskal-Wallis rank sum test, χ^2^(14) = 33.385, *p* < 0.01) but not on GOB (χ^2^(12) = 17.68, *p* = 0.12) phenotype magnitude. **C,D:** Grey dots indicate impact on binding affinity of those 1000 Genomes variants carried by the LCL samples which do not test for significant asymmetric binding (AlleleSeq [23], FDR threshold = 0.1) and are not, therefore, VDR-BVs.

To distinguish these possibilities we next compared the population frequencies of alleles associated with either historical Loss Of Binding (LOB) or Gain of Binding (GOB) affinity to VDR (Methods). LOB VDR-BVs cause significantly stronger effects when within either 5’ or 3’ hexamer than within the DR3 spacer sequence (Wilcoxon rank sum test, *p* < 0.01); GOB variants: *p* = 0.11. VDR-BVs affecting one of the most highly conserved motif positions (position 13) always result in motif-breaking LOB effect (Figure 5C and D; Supplementary Figure 18C and D).

We next found that RXR: VDR motif sequences have been under constraint during recent evolution, within the extant human population. We did so by comparing the frequencies of derived alleles (Derived Allele Frequency [DAF] test) for LOB or GOB VDR-BVs in either Northern and Western European (CEU) or Yoruban (YRI) populations. In absence of demographic confounders, stronger purifying selection is implied by an excess of low population frequency variants in one set of variants over another [39].

We detected significant increases in DAF distribution for GOB over LOB VDR-BVs, both for CEU (Figure 6C, top-left panel) and YRI (Figure 6C, bottom-left panel) cohorts. Class I binding site VDR-BVs exhibited significant differences in GOB versus LOB DAF distributions (Supplementary Figure 19; *p* < 0.05 for both CEU and YRI). By contrast, variants in RXR::VDR hexamers not assigned as VDR-BVs showed no significant difference in GOB versus LOB DAF distributions (Figure 6C, top-right and bottom-right panels). Significant differences between GOB and LOB DAFs were attained even when observing the RXR and VDR motif hexamers separately, and in both sub-populations (*p* < 0.012). Consequently, LOB variants are under substantially stronger purifying selection than are GOB variants. This is an important finding because it indicates that reduced binding of VDR at these genomic positions, and presumably reduced vitamin D levels, are commonly deleterious.

**Figure 6:**
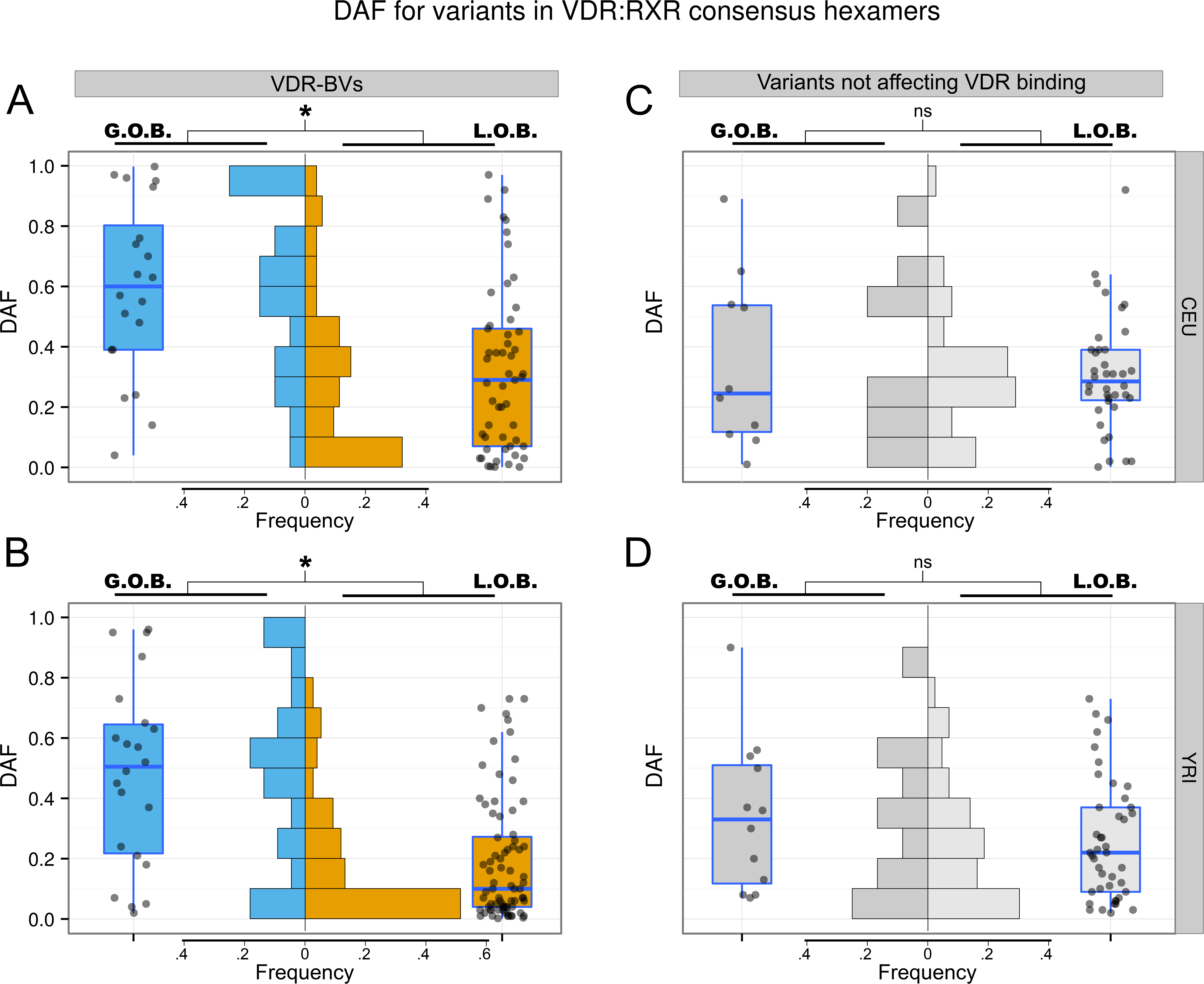
Evolutionary conservation of VDR-BVs at RXR::VDR consensus motifs. The diagrams show distributions of the Derived Allele Frequency (DAF) for variants in RXR::VDR consensus motifs within reproducible CP_*o*3_ binding peaks. DAF values are separated by ethnicity (panels **A** and **C:** CEU; panels **B** and **D:** YRI); variants are separated by their effect on VDR binding affinity (panels **A** and **B:** VDR-BVs; panels **C** and **D:** 1000 Genomes variants carried by the LCL samples which do not test for significant asymmetric binding and are not, therefore, VDR-BVs). Within each of the four panels, variants are split in two groups, based on their effect on VDR binding affinity direction (whether GOB or LOB). For all quadrants, only DAFs for variants hitting hexamer positions (i.e. hitting either the RXR recognition element at positions 1-6 or the VDR recognition element at positions 10-15) in the RXR::VDR motif are shown. An asterisk indicates significance (non parametric Wilcoxon rank sum tests,*α* = 0.05). Panel **A:** Wilcoxon rank sum test, *p* = 2.8 × 10^−4^. Panel **B:** Wilcoxon rank sum test, *p* = 4.6 × 10^−5^. Panels **C** and **D:** *p* = 0.55 and *p* = 0.14 respectively. *ns* = not significant.

## Discussion

We have demonstrated how genetic variation alters the binding affinity of the vitamin D receptor (VDR), a member of the nuclear receptor family of transcription factors. Genetic variants significantly associated with immune and inflammatory disease were found to disrupt VDR binding and to be located preferentially within enhancer regions (Figure 3), in line with a recent analysis of causal immune disease-related variants [6]. The study benefited from the ∼5-fold greater spatial resolution and signal-to-noise ratios provided by the ChIP-exo assay.

In its best studied model of allosteric modulation, the RXR::VDR heterodimer binds to DNA and enhances transcription via response elements that typically consist of two hexameric ((A/G)G(T/G)TCA) half-sites, with RXR and VDR DNA binding domains occupying the upstream and downstream half-site, respectively [40]. This DR3 motif, which we previously identified using ChIP-seq data from two calcitriol-activated CEU LCLs [14], was also the predominant VDR binding interface in calcitriol-activated LCLs in the present ChIP-exo study. We identified strong enrichment of VDR binding within enhancers, with stronger enrichments associated with the more stringent subsets of VDR-BVs (Figures 1 and 3). Whilst VDR binding occurs preferentially within promoters, these interactions are only rarely associated with RXR::VDR DR3 motifs, and may be mediated by additional DNA-binding co-factors, including CTCF, and might reflect unproductive binding to open chromatin. In contrast, those VDR binding sites containing strong instances of the RXR::VDR DR3 motif were particularly enriched in enhancers, and also near to genes involved in immunity processes and related diseases (Figure 2). VDR thus is expected to exert substantial effect on the biology of the immune system as a sequence-specific enhancer regulator.

Most remarkably, we found variants weakening or strengthening the RXR::VDR PWM score to be highly predictive of the loss or gain of VDR binding affinity (Figure 5). Furthermore, the VDR-bound RXR::VDR motif shows evidence of levels of sequence conservation across vertebrate evolution that mirror its information content, and loss of VDR-binding variants over modern human evolution has occurred under stronger purifying selection than gain of binding variants (Figure 6). Together these results imply that VDR-binding and thus presumably vitamin D levels, have in general been protective of disease.

Our study will have underestimated the true extent of regulatory variation at VDR binding sites. This is because the asymmetric binding analysis [23] by definition can only test variants which are heterozygous for a given LCL sample: therefore, true positive VDR-BVs will be unobserved if all LCL samples we considered are homozygous. Similarly, not all of the required three biallelic genotypes will have been present in these samples for the regression-on-genotype analysis [22, 21]. Consequently, a larger scale study would be expected to show improved power to detect VDR-BVs.

At least 90% of VDR-BVs lay outside of RXR::VDR DR3 motifs. The majority of variable binding could occur, as with other TFs [41], at weak or non-canonical binding motifs, or its affinity may alter because of genetic disruption to a cofactor binding site [42]. Another explanation is that VDR-BVs commonly exert their effect by altering the DNA conformation of regions flanking the core binding site [43]. Nevertheless, our *de novo* motif discovery identified several proteins as potential RXR::VDR cofactors, whose altered DNA-binding affinity could influence VDR binding. Over-representation of motifs in VDR peaks has been observed before, for example in a recent study of VDR binding in monocytes and monocyte-derived inflammatory and tolerogenic dendritic cells [44]. Direct interaction between VDR and CREB1 has been reported previously [45] as have functional interactions between vitamin D/VDR and oestrogen metabolism and MYC proteins [46, 47]. Co-occurrence of GABPA and ESRRB motifs with VDR-binding sites had been reported previously [48]. ZNF423 is not known to bind VDR yet it might do so indirectly because it physically associates with its heterodimer partner, RXRα [49].

Our finding that VDR-BVs are significantly enriched within eQTLs (Figure 3C) is intriguing because the eQTLs were inferred from LCLs not treated with calcitriol, and implied that these enhancers’ activities may not be entirely vitamin D-dependent in these cells. Nevertheless, there was no clear concordance or discordance between the direction of change of VDR binding at VDR-BVs (in calcitriol-treated LCLs) with the direction of change of gene expression in calcitriol-minus human LCLs reported for the GEUVADIS eQTL variants (data not shown). This, and the lack of enrichment of the DR3 motif in the VDR binding sites for the basal (unstimulated) state in LCLs [14] indicate that a calcitriol-activated LCLs’ study at a similar scale to those employing unstimulated LCLs [28] will be required to fully resolve this issue.

Our most important results derived from a conservative analysis of a stringent set of replicated binding variants (Figure 4B). From this, we report a significant excess of VDR-rBVs coincident or in strong LD with genome-wide significant GWAS tag variants for six disorders, including three that are autoimmune disorders (inflammatory bowel disease, Crohn’s disease and rheumatoid arthritis) whilst a fourth, endometriosis, is frequently comorbid with autoimmune disorders [50]. Deficiency of serum 25(OH)D levels is associated with cardiovascular disease risk factors in adults [10], while the association between vitamin D levels and coronary artery disease, as for many disorders, is debated [51]. These results associate a molecular phenomenon, the genetic disruption of nuclear receptor binding, with a narrow set of immune-related syndromes and diseases.

Based on the combined functional, evolutionary and GWAS-based evidence, we propose that VDR-BVs represent good targets for subsequent experimental validation using, for example, genome editing in cells. It is also expected that only a small minority of genes bound by VDR will be altered in expression. A large-scale expression QTL study in calcitriol-activated LCLs will thus be required to further narrow down the set of candidate disease genes whose regulatory elements are variably bound by VDR and which are differentially expressed across the human population.

We have provided the first quantitative association-based analysis to explain the genetic effect on binding affinity variation for a nuclear receptor, and to propose a mechanistic model for disease susceptibility. Our results support the hypothesis that DNA variants altering transcription factor binding at enhancers contribute to complex disease aetiology and suggest that altered VDR binding, and by inference variable vitamin D levels, explain, in part, altered autoimmune and other complex disease risk.

## Materials and Methods

### VDR ChIP-exo Sample Preparation

Calcitriol stimulated lymphoblastoid cell lines (LCLs) were grown and prepared for ChIP as described previously with some modifications [52] and adapted for ChIP-exo [15, 16]. LCLs are known to be a good model of primary B-cells [53] and we chose lines from HapMap, a unique resource whose genomes have already been sequenced. Briefly, cells were incubated in phenol red free RPMI-1640, 10% charcoal stripped FBS, 2mM glutamine-L, penicillin with streptomycin solution (100 U/mL + 100 microG/mL) medium at 37C and 5% CO_2_. Cells were harvested after stimulation for 36 hours with 0.1 *μ*M calcitriol (Sigma) and crosslinked using a 1% formaldehyde buffer for 15 minutes at room temperature and quenched with 0.125 M glycine. Cells were lysed and chromatin sheared by sonication into fragments of ∼200-1000 bp. VDR-bound genomic DNA regions were isolated using a rabbit polyclonal antibody against VDR (Santa Cruz Biotechnology, sc-1008). Immunoprecipitated chromatin was then processed enzymatically on magnetic beads. Samples were polished, A-tailed, ligated to sequencing library adaptors and then digested with lambda exonuclease to remove nucleotides from 5’ ends of double stranded DNA. Single-stranded DNA was eluted and converted to double-stranded DNA by primer annealing and extension. A second sequencing adaptor was ligated to exonuclease treated ends, PCR amplified, gel purified and sequenced. Libraries were prepared for Illumina as described in [16].

### Data Processing and Peak Calling

Full data processing steps are presented in the Supplementary Material. Briefly, we mapped the ChIP-exo sequence reads (Peconics Inc., USA; single-end 40bp) against the hg19 build of the human reference genome [54] using BWA (v. 0.7.4, [55]) followed by Stampy (v 1.0.21 [56]). We filtered the reads to retain only uniquely mapping reads with MAPQ > 20 and removed reads mapping to empirical blacklist regions identified by the ENCODE consortium [24].

For peak calling, we first computed strand cross-correlation profiles [57] of read start densities to obtain consensus estimates of the ChIP-exo digested fragment sizes, which we corrected for the presence of phantom peaks [58] using a mappability-corrected approach to cross-correlation inference (MaSC, [59]). We used the cross-correlation based estimates for the digested fragment size as a MACS2 (version 2.0.10, [60]) input parameter to obtain peak calls. Separately, to increase sensitivity, we called peaks using GPS/GEM (v. 2.4.1, [61]) with recommended ChIP-exo options `-smooth 3-mrc 20`. We then pooled, for each sample, the two sets of peak calls (Supplementary Material). Finally, we used DiffBind [62] to manipulate the per-sample merged peaksets and to obtain consensus peakset (CP_*o*3_, CP_*o*10_, CP_*o*20_) based on an overlap threshold. To do this, we normalised the read numbers using DiffBind’s embedded EdgeR routines, based on the Trimmed Means of M’s (TMM) algorithm [63].

### Interval Overlap Enrichment Analysis

We used the Genomic Association Tester [64] to perform the randomization-based interval enrichment analyses. All analyses were based on 10,000 randomisations over the chosen background, and backgrounds differed depending on the specific analysis performed. All analyses accounted for genome-wide patterns of GC content variability and, where relevant, for uniquely mappable regions (40bp reads). For the enrichment analysis of VDR-BVs in GWAS intervals and QTLs, we used an in-house bootstrapping pipeline to perform LD-filtering (to retain only LD-independent foreground and background variants) and DAF matching. Further details on annotations, background and analytical design are provided in the Supplementary Material.

### Motif Discovery

Full motif analysis steps are documented in the Supplementary Material. Briefly, we carried out *de novo* motif analysis using multiple parallel approaches: MEME-ChIP [65] from the MEME suite [66], XXmotif [67] and PScanChIP [20] from the Weeder/MoDtools suite [68]. For the MEME-ChIP and XXmotif analyses, we used the peak intervals in CP_*o*10_ and extracted the sequence underlying each interval ±150 bp from a repeat-masked version of the hg19 reference [54]. For the PScanChIP analysis we used as the input the full set of CP_*o*3_ binding regions with Jaspar PWM descriptors, to which we added the best heterodimeric RXR::VDR PWM found by XXmotif and the best monomeric VDR PWM found by DREME (MEME-ChIP package). The global background chosen for the analysis consisted of built-in PScanChIP DNase I digital genomic footprinting data for the LCL CEPH individual NA12865. For the motif analysis at the 43,332 VDR-BVs we used PScanChIP [20] with Jaspar PWM descriptors and a mixed background of promoters and LCL DNase cut sites.

### Differential Binding Analyses

We performed both allele-specific (VDR-ASB) and regression on genotype (VDR-QTL) differential binding analyses. Both methods start from analyses of read counts. For the VDR-QTL analysis, reads were normalised and covariant-corrected as detailed in the Supplementary material.

For the VDR-ASB tests, we utilised the sequence composition of ChIP-exo sequence reads overlapping heterozygous SNPs to determine the sequences originating from each allele separately [69, 70, 23] and to identify allele-specific binding events showing significant difference in the number of mapped reads between parental alleles. We employed a modified version of the AlleleSeq pipeline [23] to carry out the VDR-ASB analysis. Briefly, we tested for significant allelic imbalance among all read pile-ups intersecting a variant showing heterozygous genotype, given a null hypothesis of 50% paternal versus 50% maternal reads. We followed [23] in controlling for two major sources of bias typically encountered when running an allele-specific analysis of short read data: a bias due to mapping to the reference hg19 genome [71] and a bias due to unannotated copy number variants skewing the read counts for some of the SNPs being tested [72]. In addition, we corrected for a third source of potential erroneous ASB SNP calls by removing any significant allelic imbalance hypothesis falling under repeat regions contained in the ENCODE blacklist data [24].

For the VDR-QTL tests, we analysed all 27 LCL samples simultaneously. We grouped the samples based on the genotypes of the underlying variant, and carried out QTL association testing between the variant’s genotype (under the assumption of an additive genetic model [73]) and the phenotype at that location (the VDR binding affinity based on the quantile-normalised ChIP-exo peak size at that location). We used an in-house pipeline based on SNPs and indels imputed with IMPUTE2 [74] and Bayesian regression modelling based on SNPTEST [21] and BIMBAM [22]. Bayesian methods for analysing SNP associations are now an established tool in GWAS analysis [75, 76, 77] and show advantages over the use of *p*-values in power and interpretation [73] though they require tighter initial modelling assumptions when compared to frequentist methods. For details on the underlying model and priors we refer the reader to the Supplementary document.

### Variant Annotation

We annotated all variants associated with differences in VDR binding and obtained by pooling the VDR-QTL and the VDR-ASB variants (VDR-Binding Variants, VDR-BVs) according to the guidelines in [25] and using a customised version of the Funseq2 suite of algorithms [26]. We added additional VDR-centric information to Funseq2: VDR CP_*o*3_ binding regions, motif intervals from the PScan-ChIP analysis for the VDR:RXR Jaspar DR3 motif and for the DREME VDR monomer, and VDR dimer/monomer PWM information. Gene annotation used Funseq2’s default (Gencode, v. 16) and the HOT annotation in Funseq was filtered to only retain LCL HOT information. We ran Funseq using the `-m 2-nc` options so that all annotation referred to the ancestral allele of the variant.

### Availability of data and material

Sequencing data from this study have been submitted to the Gene Expression Omnibus archive (GEO; http://www.ncbi.nlm.nih.gov/geo; accession number GSE73254).

## Acknowledgements

We thank Mark Gerstein, Jieming Chen, Joel Rozowsky, Alexej Abyzov, Ekta Khurana and Yao Fu for precious help and technical insight on the AlleleSeq, CNVnator and Funseq2 platforms. We also thank Matthew Stephens and Yongtao Guan for help and insight on the PIMASS and BIMBAM platforms. We thank Gosia Trynka and Kamil Slowikowksi for providing *r*^2^-based LD block data; Yuchun Guo and David Gifford for precious help and technical insight on the GEM peak caller; Gerton Lunter for help on the STAMPY platform; Andreas Heger for help and technical insight on the GAT software and genomic randomization testing; Rory Stark and Simon Anders for insight on differential analysis of ChIP-seq data; Laura Clarke for help with the 1000 Genomes data and Daniel Gaffney for providing helpful pointers regarding the GEUVADIS QTLs calls; We thank Li Shen and Ning-Yi Shao (Icahn School of Medicine at Mount Sinai) for insight into NGSplot utilisation; and Oscar Bedoya Reina and Christoffer Nellaker for helpful discussions.

## Authors’ contributions

AJBT and SR designed the experiment and initiated the study; AJBT performed the ChIP-exo experiments and generated the data; GG devised the analytical strategy and undertook all computational analyses with guidance from WH, SR and CPP; GD assisted in the design of the experiment and in data generation, and revised the manuscript; GG and CPP wrote the manuscript with contributions from AJBT and SR.

**Abbreviations**
ASB: Asymmetric Binding Analysis
ChIP: Chromatin Immuno Precipitation
dsQTL: DNase I sensitivity Quantitative Trait Locus
DAF: Derived Allele Frequency
DHS: DNase I Hypersensitive Site
eQTL: expression Quantitative Trait Locus
ENCODE: Encyclopedia of DNA Elements
FDR: False Discovery Rate
GOB: Gain of Binding
GWAS: Genome Wide Association Study
HOT: High-Occupancy Target
LCL: Lymphoblastoid Cell Line
LOB: Loss of Binding
MHC: Major Histocompatibility Complex
PCR: Polymerase Chain Reaction
PWM: Positional Weight Matrix
QTL: Quantitative Trait Locus
RXR: Retinoid X Receptor
SNP: Single Nucleotide Polymorphism
TF: Transcription Factor
TFBS: Transcription Factor Binding Site
TSS: Transcriptional Start Site
TTS: Transcriptional Termination Site
VDR: Vitamin D Receptor
VDR-BV: VDR Binding Variant
VDR-rBV: VDR reproducible Binding Variant
VDR-sBV: VDR stringent Binding Variant
25(OH)D: 25-hydroxyvitamin D

